# Large-scale, quantitative protein assays on a high-throughput DNA sequencing chip

**DOI:** 10.1101/342808

**Authors:** Curtis J Layton, Peter L McMahon, William J Greenleaf

## Abstract

High-throughput DNA sequencing techniques have enabled diverse approaches for linking DNA sequence to biochemical function. In contrast, assays of protein function have substantial limitations in terms of throughput, automation, and widespread availability. We have adapted an Illumina high-throughput sequencing chip to display an immense diversity of ribosomally-translated proteins and peptides, and then carried out fluorescence-based functional assays directly on this flow cell, demonstrating that a single, widely-available high-throughput platform can perform both sequencing-by-synthesis and protein assays. We quantified the binding of the M2 anti-FLAG antibody to a library of 1.3×10^4^ variant FLAG peptides, exploring non-additive effects of combinations of mutations and discovering a “superFLAG” epitope variant. We also measured the enzymatic activity of 1.56×10^5^ molecular variants of full-length of human *O*^6^-alkylguanine-DNA alkyltransferase (SNAP-tag). This comprehensive corpus of catalytic rates linked to amino acid sequence perturbations revealed amino acid interaction networks and cooperativity, linked positive cooperativity to structural proximity, and revealed ubiquitous positively-cooperative interactions with histidine residues.

## Introduction

High-throughput DNA sequencing technologies have enabled the investigation of diverse biological processes wherein the functional consequences of nucleic acid variation can be linked directly to the abundance and sequence of DNA fragments that are quantified at scale. These applications (e.g. ChIP-seq, ATAC-seq, bisulfite sequencing, Hi-C, bind-n-seq, etc. [1–5]) largely define the contemporary methodological foundations of modern functional genomics. In contrast, methods for directly assaying the influence of *protein* sequence variation on function has remained challenging to similarly scale and disseminate [6]. *In vitro* approaches for high-throughput protein functional measurements have included the quantification of selective enrichment [7–9] of protein particles linked to their encoding nucleic-acids [10, 11] and parallelized binding assays on protein and peptide microarrays [12], including arrays generated with *in vitro* protein translation [13–15] and *in situ* peptide synthesis [16–18]. However, existing implementations have not yet provided the scalability, simplicity, automation, or accessibility necessary for widespread application. The implementation of direct and quantitative assays of protein function with the automation and throughput of a modern high-throughput sequencing platform would greatly expand our ability to develop and test a predictive understanding of the functional impact of coding mutations, to identify and characterize amino acid interaction networks and dependencies, and to learn useful principles and paradigms for rational design of protein function.

To enable facile, widely deployable, high-throughput, and quantitative protein characterization, we sought to leverage the capabilities and widespread adoption of the now-ubiquitous DNA sequencing flow cells to directly assay protein function at scale. High-throughput flow cell DNA arrays, the core of Illumina sequencing [19], have recently been repurposed for quantitative high-throughput investigation of DNA-protein, RNA-protein, and RNA-RNA interactions across nucleic-acid sequence space. Building on this and other work [9, 20–24] we have reengineered high-throughput DNA sequencing methods to assay protein function across a vast polypeptide sequence space. This approach aims to bring quantitative protein functional investigation to DNA sequencing-scale throughput using a hardware platform and microfluidic chip compatible with fluorescence-based sequencing by synthesis (SBS) methods, demonstrating that a single, widely-available high-throughput platform can, in principle, perform both sequencing-by-synthesis and protein function assays.

## Results

To develop methods capable of generating and quantifying our protein array on an SBS-compatible platform, we constructed a flexible, programmable workstation capable of microfluidically interfacing with and imaging sequencing flow cells [23, 25]. The resulting TIRF microscopy platform, based on the automated fluidics and fluorescence microscopy components of an Illumina GAIIx sequencer, interfaces with previously-sequenced (and therefore sequence- and position-indexed) Illumina MiSeq flow cell arrays [23]. To generate a protein array, we create a library of engineered DNA constructs that encode for polypeptides of interest (fig. S1), then sequence this library on an Illumina MiSeq. After moving the sequenced chip to our assay platform, we re-register the cluster positions to their sequences (see methods). We next perform *in vitro* transcription/translation on chip such that both the transcript and nascent peptide remain associated with their DNA template, producing a tethered protein array. To enable this tethered display, each member of the sequencing library contains DNA sequence elements that allow for 1) prokaryotic *in vitro* transcription, 2) immobilization of the resulting RNA transcript, 3) translation, and 4) ribosome stall, similar to ribosome display [26] (Fig 1A). Fluorescence-based functional assays (e.g. quantifying binding of fluorescently-labeled binding partners or incorporation of fluorescent substrates) may then be conducted directly on this array. Fluorescence images, paired to cluster DNA sequences generated from the sequencing run, are then quantified (see methods) to assay polypeptide binding or other protein function. We call this method **Prot**ein display on a **Ma**ssively **P**arallel Array, or Prot-MaP.

**Fig 1.**
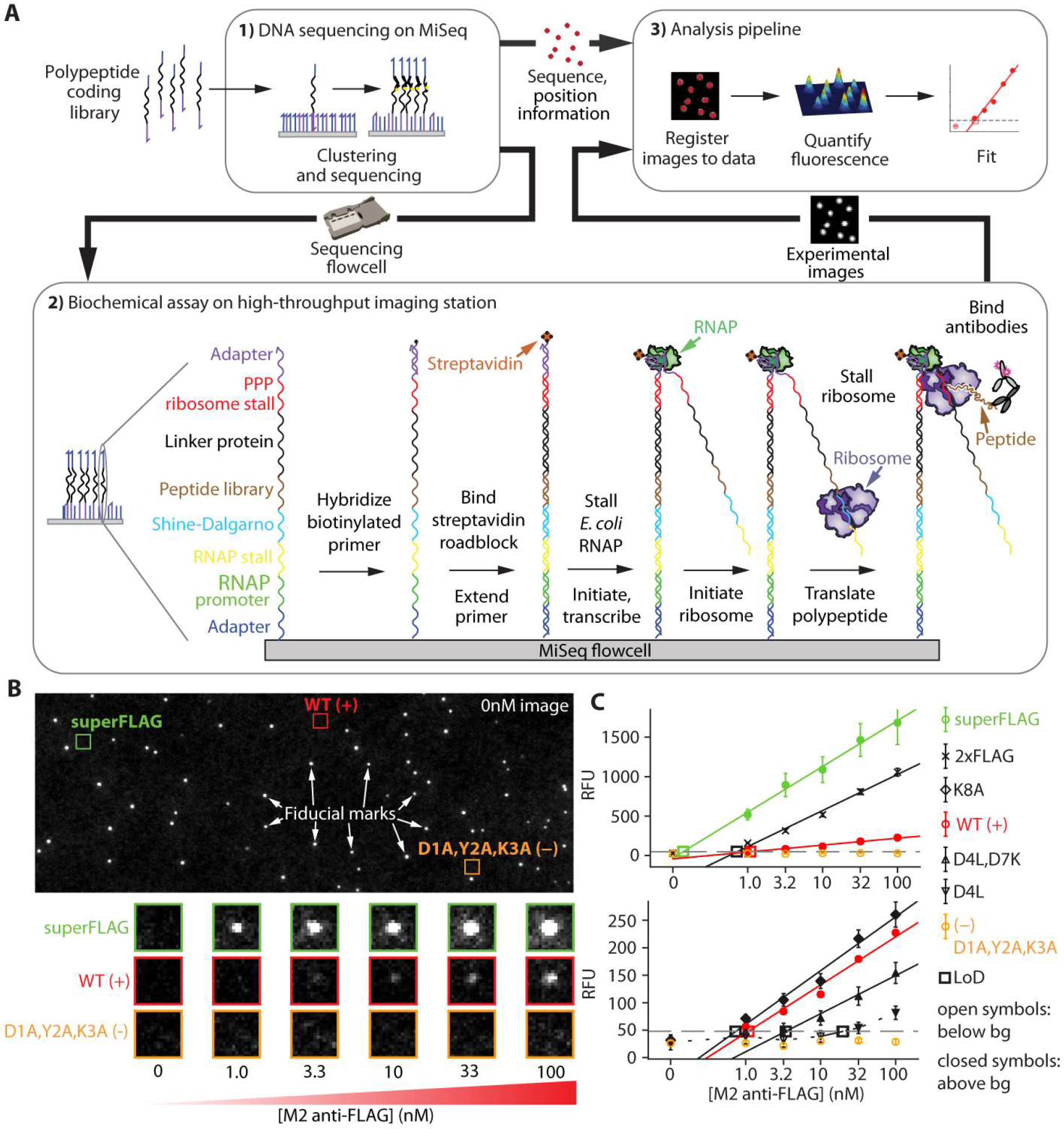
A high-throughput protein array on a DNA sequencing flow cell. (A) 1.) A polypeptide-encoding DNA library is clustered and sequenced on a MiSeq, then the chip is transferred to an imaging station where 2.) roadblocked transcription and stalled translation takes place *in vitro*, producing bound polypeptides. A primary antibody and fluorescently-labeled secondary antibody are then bound to the array and imaged. 3.) Fluorescence images are registered to the sequence information, then quantified and fit. (B) A representative flow cell image shows fiducial marks and cluster-of-interest positions before the binding assay (top). Experimental images show fluorescent secondary antibody detection of binding across increasing concentrations of M2 anti-FLAG primary antibody (bottom). (C) Quantified fluorescence medians that rise above a background threshold (grey dashed line) are extrapolated (solid) or interpolated (dashed) to estimate the limit of detection (LoD, open squares) for each FLAG variant, including WT FLAG (DYKDDDDK) and negative control (AAADDDDK) as well as superFLAG variant (DYKDEDLL), which, like 2xFLAG (DYKDDDDKGDYKDHD), gives much higher signal than WT. Two different scales are shown (top vs bottom) to accommodate the wide dynamic range of observed signals. Open symbols indicate points below background, closed symbols are above.

As a testbed for this methodology, we characterized the widely-used FLAG peptide/M2 antibody interaction. FLAG peptide is commonly used for protein labeling, purification, and as a general affinity reagent [27]. The consensus sequence profile of the linear peptide epitope of M2 has previously been characterized as DYKxxDxx based on 5 [28] or 18 [29] clones selected from random peptide libraries. We aimed to generate a library of target peptides that would comprehensively probe the contribution of residues in the FLAG peptide to the M2 antibody interaction with a much larger library of variants. We engineered a variant library of 13,154 sequences that encoded all single, double, and triple-combinations of mutant positions, with each position substituted to each of 6 different amino acids – A,K,D,S,F, and L – that represent small, positive, negative, polar, aromatic, and aliphatic substitutions, respectively. The peptide-coding mutant library was produced with microarray synthesis and assembled into a sequenceable library construct with elements enabling transcription, translation, and stable peptide display (Fig 1A, fig. S1, see methods).

After DNA sequencing, 623,075 total clusters encoding FLAG library members were produced with 12,739 of 13,154 (96.8%) programmed variants represented by 8 or more peptide clusters (i.e. nearly complete coverage of this synthetic FLAG library required only ~2.5% of capacity of the flow cell). After peptide generation, M2 antibody was introduced at increasing concentrations and allowed to bind to the peptide, with each concentration followed by detection with a fluorescently-labeled secondary antibody, then imaging (Fig 1B). After image registration and fluorescence quantification, the limit of detection (LoD) for antibody binding was determined for each variant (Fig. 1C, see methods), representing the lowest concentration of antibody that produces detectable binding. This Prot-MaP LoD assay is analogous to a massively multiplexed ELISA whereby primary antibody interactions are probed by secondary detection steps that produce measurable signal.

The single and double mutant affinity landscapes (Fig. 2A-B) of the canonical sequence (DYKDDDDK, “WT”) largely recapitulate the previously reported motif pattern, DYKxxDxx [28, 29]. However, we observe substantial additional constraint at position D4, with 5 of 6 mutations exhibiting no detectable binding. To investigate mutational constraint at this position further, we asked if detrimental mutations at position 4 could be rescued by mutations at other positions in our higher order mutants. Looking at all double mutants of the only measurable D4 mutant (D4L; triple mutants from WT; Fig. 2C), we observe that several combinations of mutations that include D5E and D7K partially rescue D4L, lowering LoD to near-WT levels. Many of these compensating combinations exhibit significant positive cooperativity (e.g. D4L,D5E,D7K), demonstrating the ability of Prot-MaP not only to identify critical residues and motifs, but also to establish cooperative interaction landscapes that can substantially deviate from simple additive models (Fig 2D).

**Fig 2.**
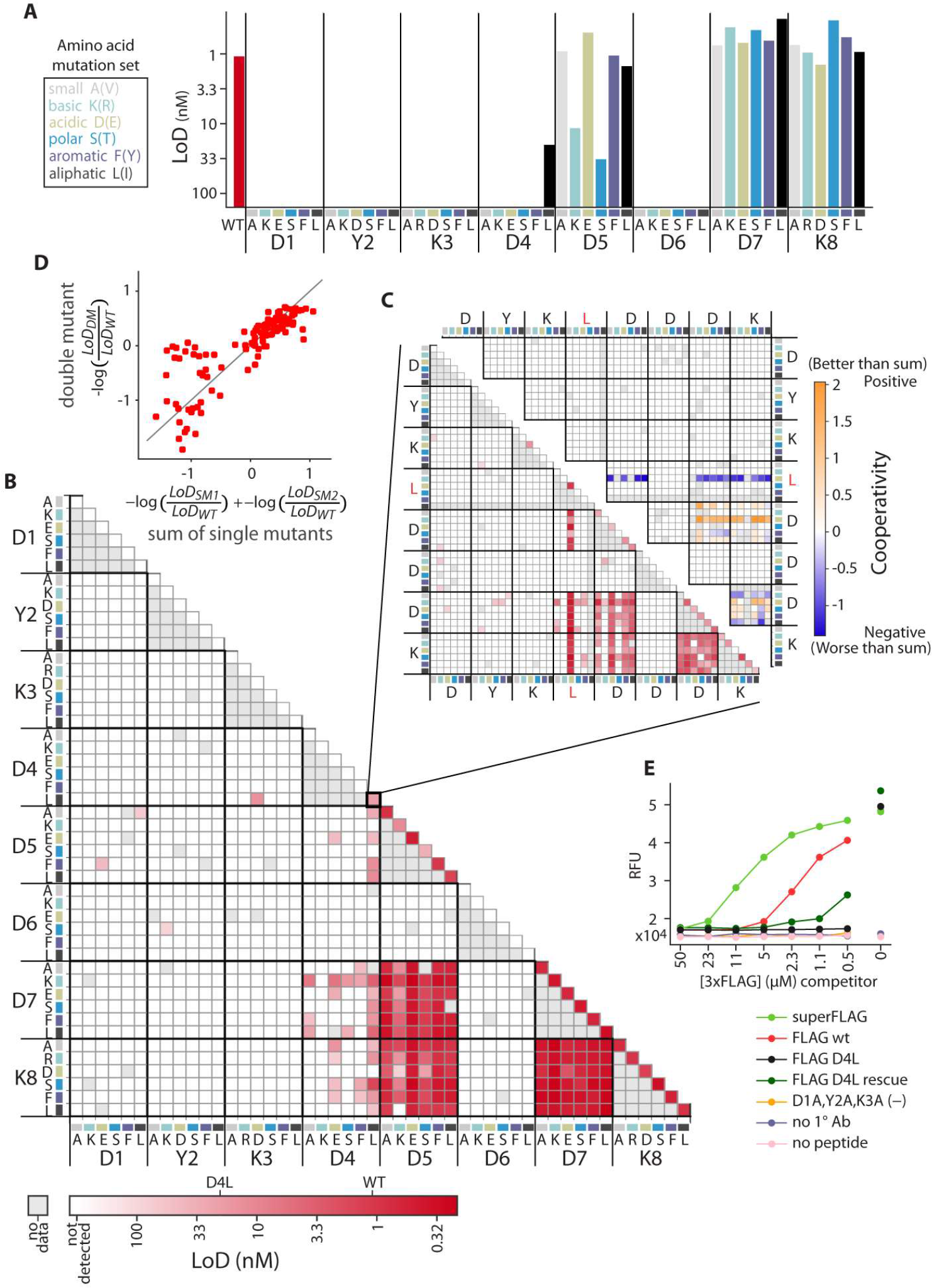
Binding landscape of FLAG peptide variants to M2 monoclonal antibody. (A) 6 amino acid mutations, each representing a physicochemical category, are represented by color. LoD for M2 anti-FLAG antibody binding to single mutants and (B) double mutants of the canonical (WT) peptide, DYKDDDDK, across all six amino acids at each position (if the WT identity is the same, the mutation in parenthesis is made). (C) LoD (bottom left) and cooperativity (top right) of double mutants of the D4L base mutant (triple mutants from the WT). Cooperativity = −log(LoD_DM_/ LoD_WT_) – (−log(LoD_SM1_/ LoD_WT_) + −log(LoD_SM2_/ LoD_WT_)), where DM refers to the double mutant and SM1 and SM2 to the two corresponding single mutants. (D) Double mutant cycles. Deviations from the y=x line (grey) indicate non-additivity (cooperativity) (E) Orthogonal investigation of individual peptide variants with a fluorescence-based plate assay. Antibodies bound to variant peptides (see legend) were challenged for 2 hours with varying concentrations of 3xFLAG competitor peptide. Fluorescence values report residual bound M2 antibody.

Interestingly, we also found several FLAG variant sequences that exhibited significantly lower LoD than WT, including a variant we term “superFLAG” (DYKDEDLL) with an extrapolated LoD of 0.14 nM, 7.9-fold lower than WT (Fig. 1C). To confirm these observations, we performed a fluorescence-based plate assay to measure M2 binding competition between immobilized variant peptides and solutions of 3xFLAG high-affinity peptide (Fig 2E). SuperFLAG displays higher avidity to the M2 antibody then all other assayed peptides, including the original DYKDDDDK immunogen to which M2 was raised [30].

Unlike chemically synthesized peptide arrays, our *in vitro* transcribed and translated array enables the possibility of generating full-length protein features [13–15]. To demonstrate this capability, we investigated the mutational landscape of SNAP-tag, an engineered version of *O*^6^ - alkylguanine-DNA alkyltransferase, a 181 AA, 20kD human DNA repair enzyme [31]. The SNAP-tag protein transfers a benzyl group from a derivatized benzylguanine substrate (often substituted with a fluorophore) to its own Cys 145, thereby covalently labeling itself. The simplicity and specificity of this fluorogenic self-labeling reaction has led SNAP-tag to become widely used as a protein labeling tag, and here it provides an elegant model system for investigation of the sequence-function relationship of an enzyme with Prot-MaP (Fig 3A).

**Fig 3.**
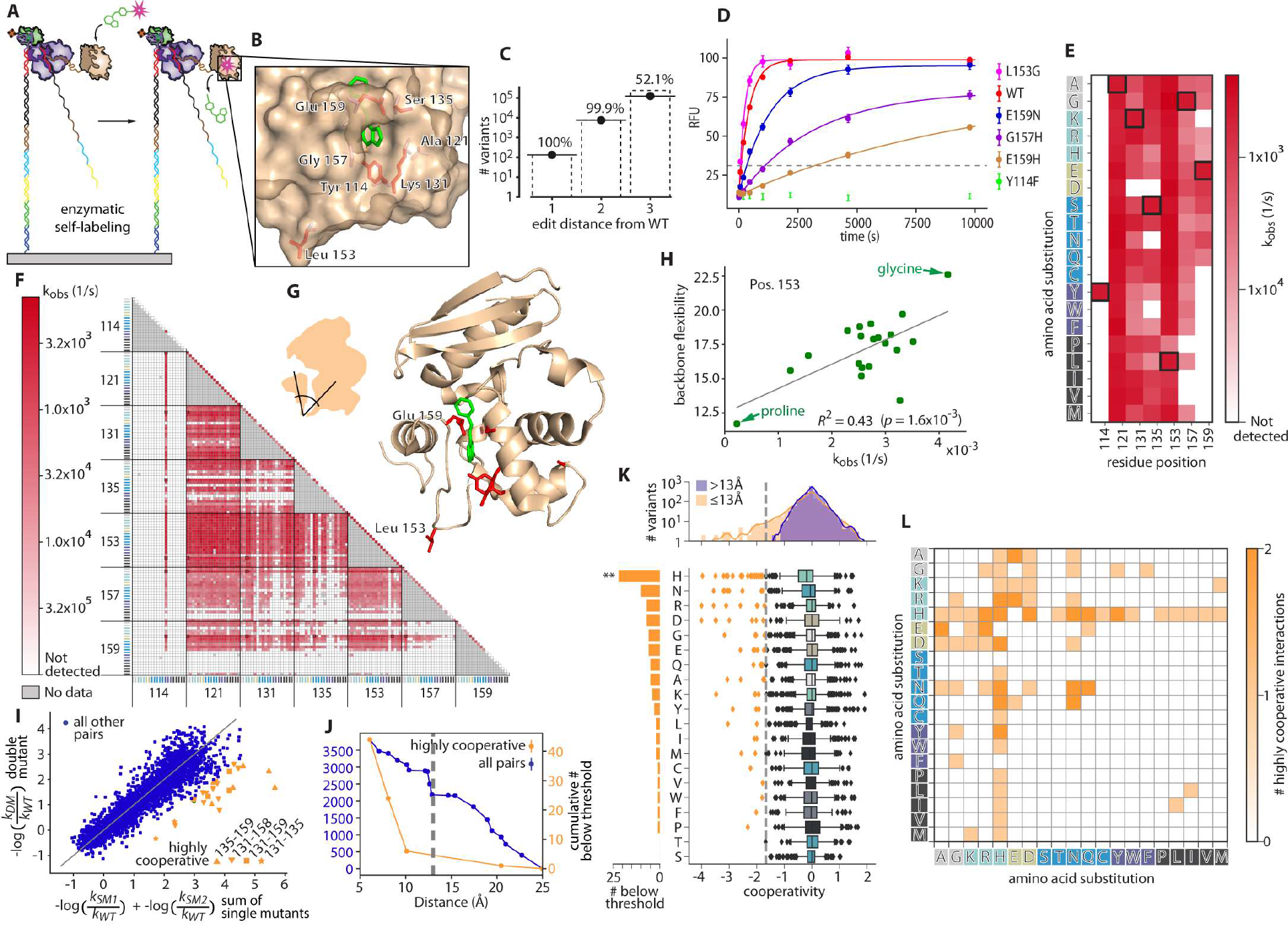
High-throughput, on-chip characterization of activity across the mutational landscape of a full-length enzyme. (A) On-chip SNAP-tag-catalyzed fluorescent self-labeling (B) 7 target amino acid positions (red) and benzylguanine substrate (green) on the structure of SNAP-tag (PDB ID: 3KZZ) (C) Library coverage of all 19 possible amino acid substitutions in single, double, and triple-combinations of the target positions. (D) Quantified fluorescence medians for selected single mutants. Variants that rose above the background threshold (grey dashed) were fit with a single exponential. (E) Fit k_obs_ values for single mutants and (F) double mutants across the 20 amino acids. WT (unmutated) variants are boxed in black. (G) Position 153 is in a loop connecting two domains. (H) Correlation of activity with backbone flexibility [34] for all 19 amino acid substitutions at position 153. R^2^=0.43 (p-value = 1.6×10^−3^ by permutations). Proline is the least flexible and glycine the most. (I) Comparison of double mutant effects on activity (y-axis) with the sum of single mutant effects (x-axis). Only a few residue position pairs (orange) demonstrate high positive cooperativity (“highly cooperative” refers hereafter to cooperativity <-1.69 (see 3K) where cooperativity = −log(k_obs,DM_/ k_obs,WT_) – (−log(k_obs,SM1_/ k_obs,WT_) + −log(k_obs,SM2_/ k_obs,WT_))). (J) Cumulative number of pairs with Cα-Cα distance within a given distance for highly cooperative (orange), and all target residue pairs (blue). Most highly cooperative interactions occur within 13Å (grey dashed) (K) cooperativity values for all target residue pairs with C_α_-C_α_ distance < 13Å (orange) and > 13Å (blue) (top) and by amino acid (counted for either or both partners) (bottom). Sum of highly cooperative pairs (grey dashed) per amino acid (left). (L) Number of highly cooperative interactions across all amino acid pairs.

To explore the relationship between protein sequence variation and catalysis in high throughput, we examined the functional interrelationship of seven residues previously identified to modulate function without entirely abolishing catalytic activity (Y114, A121, K131, S135, L153, G157, and E159) [32] (Fig. 3B). The DNA sequence encoding this mutated region (residues 114-159, the “mutant region”), was combinatorially assembled from oligos with the aim of generating single-, double-, and triple-mutant combinations of the 7 target positions across all possible 20 amino acid substitutions (fig. S1). Once assembled, the DNA fragment population was bottlenecked (by diluting to a target number of molecules) and reamplified with PCR to obtain multiple identical copies of molecular variants on the array.

After sequencing this library, RNA synthesis and *in situ* protein generation was performed on the MiSeq flow cell. We then introduced a solution of 200 nM substrate (SNAP-Surface 549) continuously onto the resulting protein array, then measured substrate incorporation over 160 minutes (see methods). A total of 6,751,654 clusters were successfully quantified across all timepoints, and signals were averaged across clusters displaying identical protein variants (> 8 clusters per variant). A total of 156,140 unique variants were assayed, including all possible single mutants across all 20 amino acids (133), 7570 of 7581 possible double mutants, 125,076 of 240,065 possible triple mutants (Fig. 3C), as well as 23,360 other variants. The majority of mutants (118,025/156,140; 75.59%) exhibited no detectable catalytic activity above background levels, while 32,072 variants exhibited detectable activity that could be well-fit by a single exponential to obtain k_obs_ (Fig. 3D, see methods).

We observed large variation in the mutational constraint for each of the 7 targeted amino acids. At one extreme, Y114 is fully constrained to tyrosine across all single and double mutants (Fig. 3E, 3F). Even Y114F, a conservative substitution that deletes only a hydroxyl moiety, is inactive [33] (Fig. 3D), suggesting that the hydrogen bonds that Y114 forms with the benzylguanine substrate are absolutely required for function. While E159 also forms a hydrogen bond with the substrate, a number of mutations to E159 retain measurable activity, suggesting that less stringent physicochemical requirements on substrate interactions at this position.

In contrast to these constrained residues, A121 is tolerant to all single mutations. L153 is tolerant to nearly all substitutions, except for proline, which is rotationally constrained in psi and phi Ramachandran angles. By exploring our catalogue of double mutants, we observed that proline is consistently the most detrimental of all the 20 amino acids at position 153 across 119/120 other single mutant backgrounds. These observations led to the hypothesis that backbone flexibility at position 153 (which is in the “hinge” region of the loop connecting helix-loop domain 153-176, including the critical residue E159, with the rest of the protein) may regulate activity via inter-domain geometry and/or dynamics (Fig. 3G). We investigated this possibility by examining the relationship between the observed enzymatic activity and amino acid backbone flexibility [34] of amino acid substitutions at position 153. We observed a strong relationship (R^2^=0.43; p-value=1.6×10^−3^; Fig. 3H), supporting this hypothesis and highlighting the utility of comprehensive mutational substitutions for testing mechanistic hypotheses.

We next aimed to characterize cooperativity in the interactions between the AAs we perturbed by analyzing double mutant cycles found within our mutational dataset (Fig. 3I). We observe a strong enrichment for physical proximity between highly positively cooperative amino acid pairs, with virtually all strong positively-cooperative interactions occurring at C_α_-C_α_ distances less than 13 Å (Fig. 3J). For example, many of the most positively-cooperative double mutants are between positions 135 and 159 – two AAs that directly physically interact in the crystal structure (PDB ID: 3KZZ). The most cooperative pair is S135D-E159R, a double mutant that produces a favorable charge-based sidechain interaction.

Interestingly, histidine appears particularly amenable to cooperative interactions. Across all positional combinations, histidine was far more likely to form a highly-cooperative interaction than any other amino acid (Fig. 3K), and did so fairly uniformly with all possible amino acid partners (Fig. 3L). Histidine can be either positively charged or neutral in different contexts, can have aromatic character, and can function as both a hydrogen bonding donor and acceptor. We hypothesize that histidine may thus function as a “Swiss army knife,” pairing promiscuously with diverse partners to produce favorable interactions and positive cooperativity. We anticipate that further investigation with high-throughput methods will provide the opportunity to explore this hypothesis in a variety of protein backgrounds.

## Discussion

Prot-MaP enables the generation of immense mutational datasets for both peptides and full-length functional proteins directly on a sequencing flow cell, allowing high-throughput analysis of mutational effects based on direct biophysical observations of protein function. Large-scale quantitative measurements of peptide-protein interaction can demonstrate and quantify nonadditivity in affinity landscapes and allow the identification of enhanced-affinity interactions (such as superFLAG). Large-scale functional analysis of full-length proteins allows data-driven characterization of functional networks and cooperative interactions of individual amino acids, geometric constraints that enforce amino acid preferences, as well as global observations and hypothesis testing regarding the functionality of specific amino acid species (e.g. histidine’s observed capacity for widespread cooperative interactions).

ProtMAP does have limitations, including a practical limit of about 1-2 kb on the length of protein-encoding DNA constructs that can be efficiently clustered and sequenced. These limits are inherited from the underlying sequencing platform and are not intrinsic to the display method. Additionally, many proteins will require different *in vitro* expression conditions and components for efficient production of functional protein (including post-translational modifications). Finally, while binding studies are broadly compatible with fluorescence-based assays, other assays (e.g. for arbitrary enzymatic activities, protein folding, or conformational changes) may necessitate the development of fluorescence-compatible measurement strategies. However, analogous to the many methods built on the foundations of high-throughput sequencing [35], we anticipate that the compatibility of Prot-MaP with widely-available high-throughput SBS platforms may serve as a catalyst for further development of specialized applications.

The implementation of facile high-throughput protein functional analysis on a broadly available sequencing flow cell demonstrates a conceptually straightforward path toward disseminated high-throughput protein functional assays using automated fluorescence imaging hardware currently used in sequencing by synthesis methods. Given the demonstrated power of high-throughput DNA sequencing methods (as applied through functional genomic methods such as ChIP-seq, Hi-C etc.) to provide insights into the relationship between DNA sequence and regulatory function, we anticipate that a similarly ubiquitous platform for high-throughput protein assays could have analogous impact on our ability to dissect biologically relevant relationships between protein sequence, structure, and function.

## Acknowledgements

We thank Gavin Sherlock, Sarah Denny, Winston Becker, and Sandy Klemm for critical reading of the manuscript. **Funding:** This work was supported by National Institutes of Health (NIH) Grant R01-GM111990 and a Technology Development Grant by the Beckman Foundation.

## Author contributions

Conceptualization C.J.L. and W.J.G.; Methodology: C.J.L. and W.J.G.; Software: C.J.L. and P.L.M.; Investigation: C.J.L.; Writing - Original Draft: C.J.L. and W.J.G.; Writing - Review & Editing: C.J.L. and W.J.G.; Visualization: C.J.L.; Funding Acquisition: W.J.G.

## Supplementary Figures

see Supplemental Information Fig. S1-S4

